# Skin Resident T Cell Interactions with NPY1R^+^ Neurons During Wound Repair Are Impaired by Obesity

**DOI:** 10.1101/2025.06.06.658336

**Authors:** Kendra Lam, Joseph Colin, Berenice Almaguer, Patricia Tulloch, Julie Jameson

**Affiliations:** Department of Biological Sciences, California State University San Marcos, San Marcos, CA 92096

## Abstract

Wound repair involves complex cellular interactions to induce efficient healing. Skin resident γδ T cells regulate keratinocyte function and inflammatory responses in wound healing through the secretion of growth factors and cytokines. Some of these cytokines are involved in the neuroimmune axis, so we aimed to ascertain if γδ T cells and neurons exhibit crosstalk during wound repair. We identified a prevalent NPY1R^+^ peripheral neuron subset in the skin. γδ T cells interact with NPY1R^+^ neurons in the epidermis and dermis of both nonwounded and wounded murine skin. To determine the impact of the γδ TCR on NPY1R^+^ neuron interactions in wound healing, we compared epidermal T cell-neuron interactions in TCRδ^−/−^ mice during wound repair. We found a decrease in interactions between epidermal αβ T cells and NPY1R^+^ neurons in TCRδ^−/−^ mice, suggesting γδ T cells communicate with NPY1R^+^ neurons more frequently than αβ T cells in the epidermis, regardless of wound repair. In contrast, dermal T cell-neuron interactions in wildtype and TCRδ^−/−^ mice during wound repair were similar between αβ T cells and γδ T cells, suggesting both T cell types mediate communications with neurons in the dermis. In obese mice there are fewer NPY1R^+^ neurons and diminished neuron-T cell interactions indicative of neuropathy. Together these findings elucidate a neuroimmune crosstalk during wound repair that becomes disrupted in obesity.

## Introduction

Wound healing is a tightly regulated process requiring communication between resident and infiltrating cells to sense damage and regulate repair (1). In the murine skin, γδ T cells monitor damage through cellular crosstalk (2). Epidermal γδ T cells respond to damaged keratinocytes and produce IGF-1, FGF-7, FGF-10, and IL-2 to induce keratinocyte and T cell proliferation, survival, and migration (3–5). Epidermal γδ T cells also regulate macrophages through cytokine and chemokine secretion (TNF-α, MCP-1) and hyaluronan deposition resulting in macrophage activation, chemotaxis, and polarization modulating the wound inflammatory state (6–8). Dermal γδ T cells are major producers of IL-17A/F, which acts through HIF-1α to induce wound reepithelialization (9). Skin γδ T cell sensing occurs early in wound healing; however, little is known about skin-resident γδ T cell and neuron interactions.

Peripheral nerve damage from injury drives a rush of immune cells, with T cells infiltrating within one week of injury (10). In the dorsal root ganglion, surgical injury of peripheral nerves induces CD4^+^ T cells to infiltrate the site, while viral infections induce CD8^+^ T cells to gather at the site (11, 12). Neuropeptide Y (NPY) is among the most abundant neuropeptides in the mammalian nervous system, playing a critical role in a wide array of physiological processes (13). Among the NPY receptors, Neuropeptide Y receptor type 1 (NPY1R) is expressed by murine dendritic cells, natural killer cells, mast cells, T cells, B cells, and macrophages (14). NPY and NPY1R can exhibit bimodal effects on the immune system, such as negatively regulating T cells and activating antigen presenting cells (14). Painful inflammation induces NPY1R upregulation in dorsal root ganglia neurons while NPY is unaffected; additionally, NPY and NPY1R have increased expression in the dorsal horn (15, 16).

Innervation within the skin depends on the integrity of the tissue itself and can be impacted by factors such as inflammation, tissue damage, and stress (17). Diabetes induces peripheral nerve damage associated with pain, numbness, and/or weakness, which compromises skin innervation (18). Type 2 diabetes and obesity are often linked to neuropathy, with approximately 60% of all diabetic patients having diabetic neuropathy (18, 19). Type 2 diabetes is associated with several other alterations to skin barrier function such as increased epidermal permeability and delayed barrier recovery, which can elicit and aggravate inflammation (20). Additionally, the chronic elevation of TNFα associated with obesity and type 2 diabetes can further drive inflammation within the skin (21). As a result of an inflammatory environment, skin γδ T cells are rendered unresponsive to stimulation by keratinocytes and lose the ability to secrete cytokines (21). However, the effects of obesity and type 2 diabetes on the neuroimmune interactions between skin γδ T cells and sensory neurons are unknown.

In this study, we investigate the interactions between skin resident T cells and NPY1R^+^ neurons as well as NPY1R expression in T cells during wound healing and obesity-related skin complications. We identify a subset of NPY1R^+^ neurons in both WT and TCRδ^−/−^ mice, independent of wound status. NPY1R expression is also observed on dermal γδ and αβ T cells, with fewer dermal αβ T cells expressing NPY1R, suggesting a distinct role for dermal γδ T cells in NPY1R-mediated functions. Notable differences emerge between epidermal and dermal γδ T cell interactions with NPY1R^+^ neurons: epidermal γδ T cells exhibit significantly higher interaction rates with NPY1R^+^ neurons than αβ T cells, while dermal γδ T cells do not show this specificity. Furthermore, diabetic and obese conditions lead to reduced NPY1R^+^ neurons and diminished neuron-T cell interactions in the dermis, highlighting potential impacts of metabolic conditions on neuronal-immune interactions in the skin.

## Materials and Methods

### Mice

Female wild-type C57BL/6J (B6) and TCRδ^−/−^ mice were purchased from The Jackson Laboratory (Bar Harbor, ME) and housed at California State University San Marcos (CSUSM). Male wildtype C57BL/6NTac mice were purchased from Taconic Biosciences (Rensselaer, NY) at 20 weeks of age after being maintained on either a 60 kcal% high-fat diet (HFD) or a 5 kcal% low fat diet (LFD) for 14 weeks. Upon arrival at CSUSM, they continued their respective HFD (Research Diets, NJ) or LFD (Zeigler Wafer, PA) for 3 additional weeks, totaling 17 weeks on the assigned diet. All mice were periodically weighed, and blood glucose monitored by an Ascensia Elite XL blood glucose monitor (Bayer, GER). All mice were studied at 12 to 24 weeks of age. Mice were given access to food and water ad libitum. All experimental procedures involving animals were reviewed and approved by the Institutional Animal Care and Use Committee of California State University San Marcos (21–003).

### Wounding Model

Mice fed a HFD or LFD were anesthetized with a mixture of 2.5% isoflurane and 1.75L/m O_2_. Full-thickness, 2-mm biopsy punch wounds were performed on the upper dorsal surface of mice. Full-thickness, 2-mm biopsy punch wounds were performed on 1 ear of the mice. Mice were housed individually and monitored daily. Mice were euthanized 7 days post wounding for dorsal wounds and 10 days post wounding for ear wounds.

### Epidermal Sheet Immunofluorescent Staining and Microscopy

Epidermal sheets were prepared as previously described (22). Briefly, Nair^TM^ was applied to ears for hair removal and washed off using DI water. Ears were separated in half and floated dermis-side down on 3.6% ammonium thiocyanate in dPBS for 15-20 minutes at 37°C with 5.0% CO_2_. Epidermal sheets were separated from the dermis, fixed with acetone for 10 minutes at RT, and stained with 4µg/mL anti-Vγ5 (Clone 536, BioLegend, San Diego, California), 4µg/mL anti-CD3ε (Clone 145-2C11, BioLegend), and/or 4µg/mL anti-NPY1R (Clone 3A1, USBiological, Salem, Massachusetts) in PBS for 1 hour at 37°C with 5.0% CO_2_. Epidermal sheets were incubated in 1xPBS for 5 minutes and mounted with SlowFade Gold Antifade Mounting Media with DAPI (Invitrogen by Thermo Fisher Scientific, Waltham, Massachusetts). The slides were viewed using an immunofluorescent microscope (Nikon DS-Qi2t, Nikon, Tokyo, Japan). Images were acquired at original magnification x400 and then analyzed using Photoshop CS2 software (Adobe, San Jose, California). Interactions between NPY1R and γδ T cells were quantified as visual overlap in B6 mice and interactions between NPY1R and αβ T cells were quantified in TCRδ^−/−^ mice. At least 12 regions were imaged per epidermal sheet per ear, accounting for both wounded and non-wounded samples. A total of 18 epidermal sheets and 669 images were captured.

### Frozen Sample Preparation and Immunofluorescent Staining

Whole dorsal skin tissue was shaved and cut into 1 cm by 2 cm sections (6 for non-wounded samples and 2 for wounded samples). The sections were then placed midline down in cryomolds and frozen using Optimal Cutting Temperature (OCT) compound on dry ice. Tissue blocks were subsequently stored at −80°C. To obtain skin sections for immunofluorescent microscopy, 10 µm skin sections were cut using a Leica Cryostat (Leica Biosystems, Nussloch, Germany). Sections were fixed with 4% paraformaldehyde for 10 minutes and incubated with blocking solution (2.5% Normal Goat Serum, 2.5% Normal Donkey Serum, 1% BSA, 2% fish gelatin, 0.1% Triton X, 0.3g glycine) for 30 minutes at RT. Skin sections were stained for 1 hour with 4µg/mL anti-TCRγδ (Clone GL3, BioLegend) 4µg/mL anti-CD3ε (Clone 145-2C11, BioLegend), 5µg/mL anti-β3-tubulin (Clone AA10, BioLegend), 4µg/mL anti-NPY1R (Clone 3A1, USBiological) at 37°C with 5.0% CO_2_. Sections were mounted with SlowFade Gold Antifade Mounting Media with DAPI (Thermo Fisher Scientific) and viewed using an immunofluorescent microscope (Nikon DS-Qi2t, Nikon). Images were acquired at original magnification x200 and analyzed using Photoshop CS2 software (Adobe). The number of T cells, number of T cells expressing NPY1R and number of T cells interacting with NPY1R^+^ neurons were quantified per mm^2^. For wildtype (WT) and TCRδ^−/−^ mice, a minimum of 24 regions were captured per skin section, accounting for both non-wounded and wounded samples. A total of 58 skin sections and 1418 images were captured. For lean and obese mice, a minimum of 22 regions were captured per skin section, accounting for both non-wounded and wounded samples. A total of 49 skin sections and 1069 images were captured.

### Statistical Analysis

Statistical analysis was performed using GraphPad Prism (version 10; GraphPad Software, Dotmatics, Boston, MA). A two-way ANOVA for multiple group comparisons was used to determine the significance of comparisons between non-wounded (NW) and wounded (W) mice in WT, TCRδ^−/−^, and DIO conditions. All findings were considered significant at *p* < 0.05.

## Results

### NPY1R is expressed by neurons in the epidermis and dermis

NPY1R is expressed by spinal neurons to inhibit neuropathic pain upon reception of NPY (23). However, NPY1R expression by neurons in the skin is less studied. To determine if neurons express NPY1R in the epidermis and dermis, we examined NPY1R and β-3-tubulin in nonwounded and 7 day wounded WT mice. NPY1R and β-3-tubulin colocalize in the majority of cells in both the epidermis and dermis, showing that most neurons express NPY1R in the skin of WT mice (Fig. 1). Further, NPY1R expression by β-3-tubulin expressing neurons in the epidermis and dermis continues during wound healing as neurons in WT mice express NPY1R in day 7 wounds (Fig. 1).

**Figure 1.**
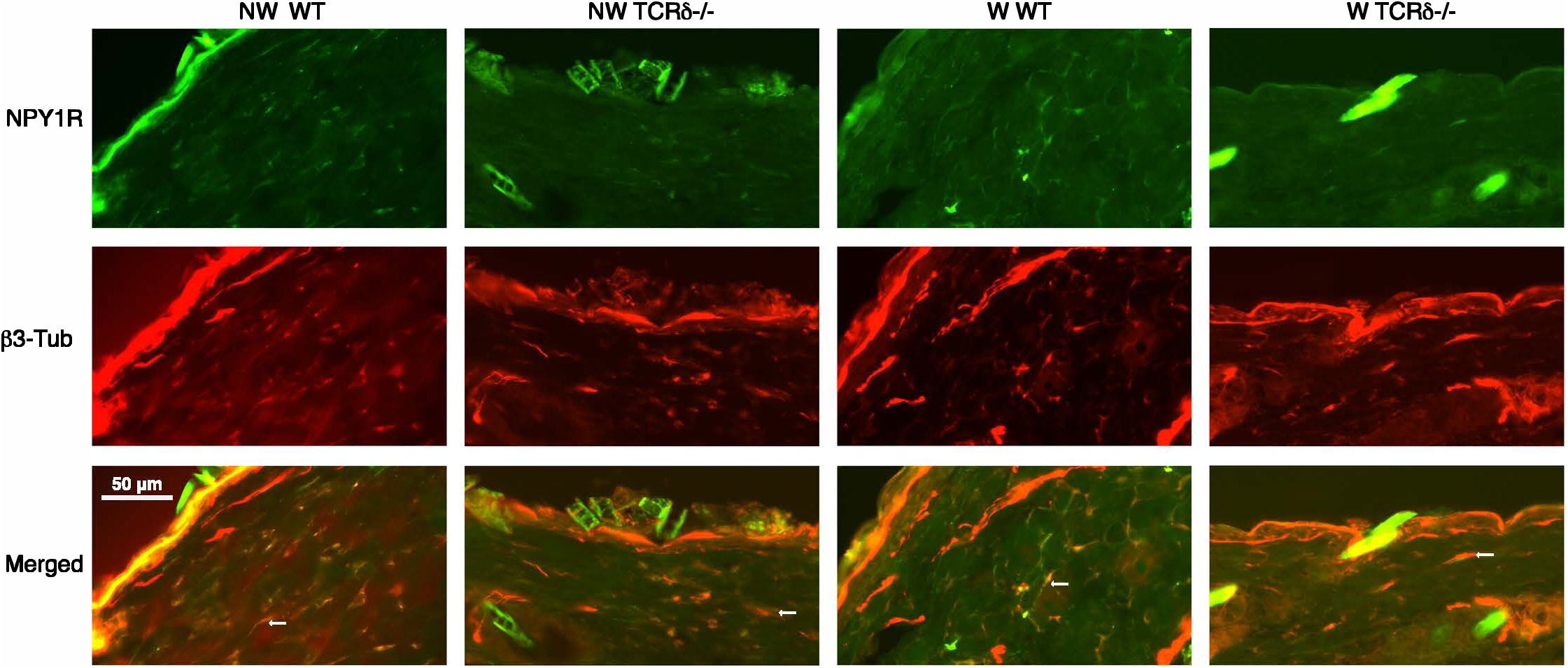
Neurons in the epidermis and dermis express NPY1R. Colocalization of NPY1R positive cells and β-3-tubulin staining show that neurons express NPY1R. Non-wounded (NW) and 7 day wounded (W) skin sections were stained with anti-NPY1R (green) and anti-β-3-tubulin (red) in wildtype and TCRδ^−/−^ mice. Images were acquired at 200x magnification.

### NPY1R expressing neurons are less prevalent in the skin of mice lacking γδ T cells

Given that γδ T cells are located in both the epidermis and dermis, where they are in close proximity to neurons and play a critical role in peripheral inflammation, we investigated whether γδ T cells have a specific role in modulating neuronal NPY1R expression during wound repair. TCRδ^−/−^ mice exhibit β-3-tubulin^+^ neurons in the epidermis and dermis confirming previous accounts showing that γδ T cells are not required for proper innervation of the skin (24) (Fig. 1). However, there was less NPY1R staining in the epidermis of TCRδ^−/−^ mice (Fig. 2A) and dimmer NPY1R^+^ staining in the dermis of TCRδ^−/−^ mice (Fig. 3A), suggesting γδ T cells may have an impact on NPY1R expression by neurons.

**Figure 2.**
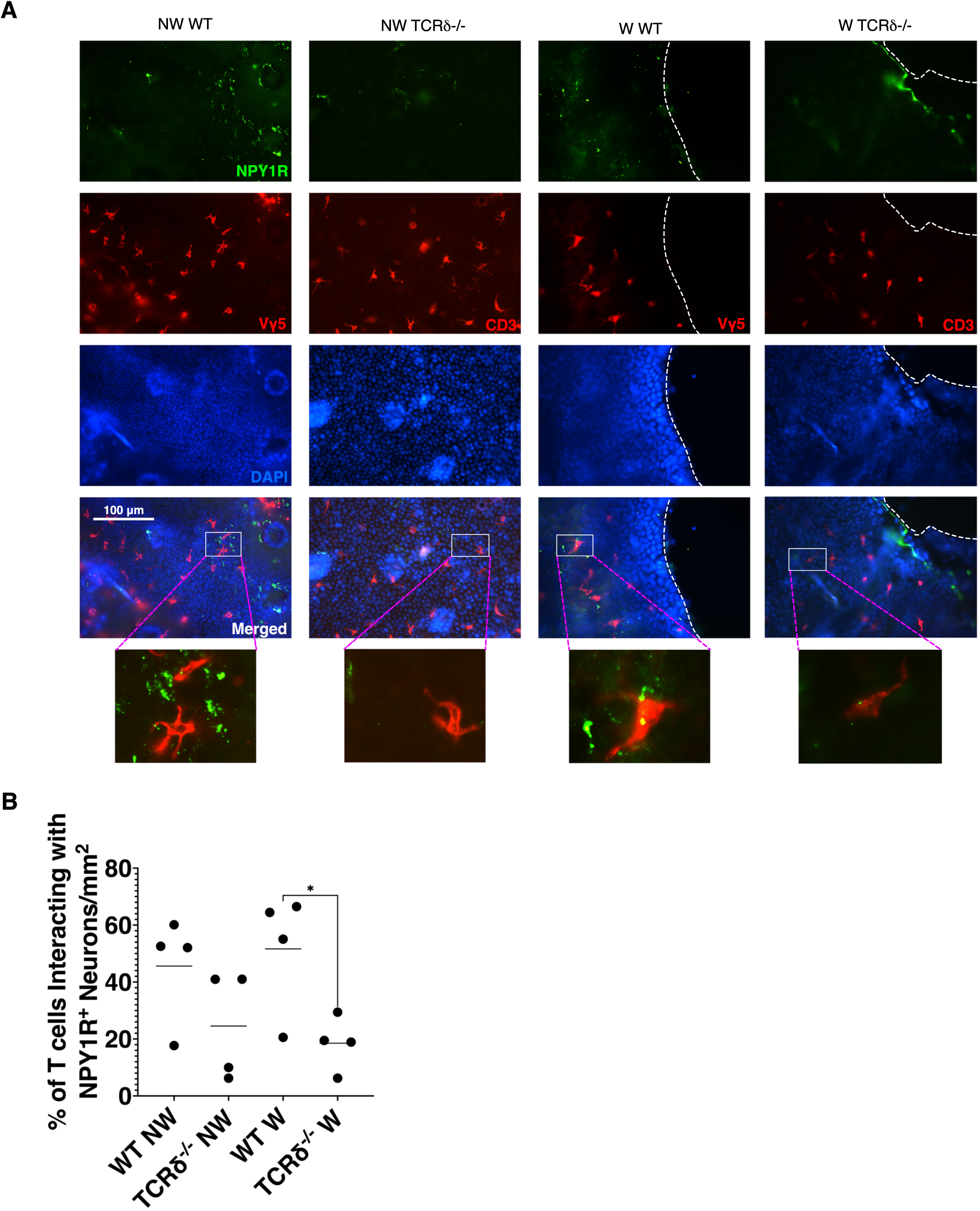
γδ T cells have a specialized role in mediating NPY1R^+^ neuron and T cell interactions in the epidermis. **(A)** Non-wounded (NW) and wounded (W) (10 day) epidermal sheets from WT and TCRδ^−/−^ mice were stained with anti-NPY1R FITC (green), Vγ5 PE (red) or CD3 PE (red), and DAPI (blue). (**B**) There are fewer interactions between T cells and NPY1R^+^ neurons in wounded mice that lack γδ T cells. Interactions between T cells and NPY1R^+^ neurons were quantified based on fluorescent overlap using immunofluorescence microscopy and Adobe Photoshop (magnification 400x). Two-way ANOVA was used for statistical analysis. *p < 0.05

**Figure 3.**
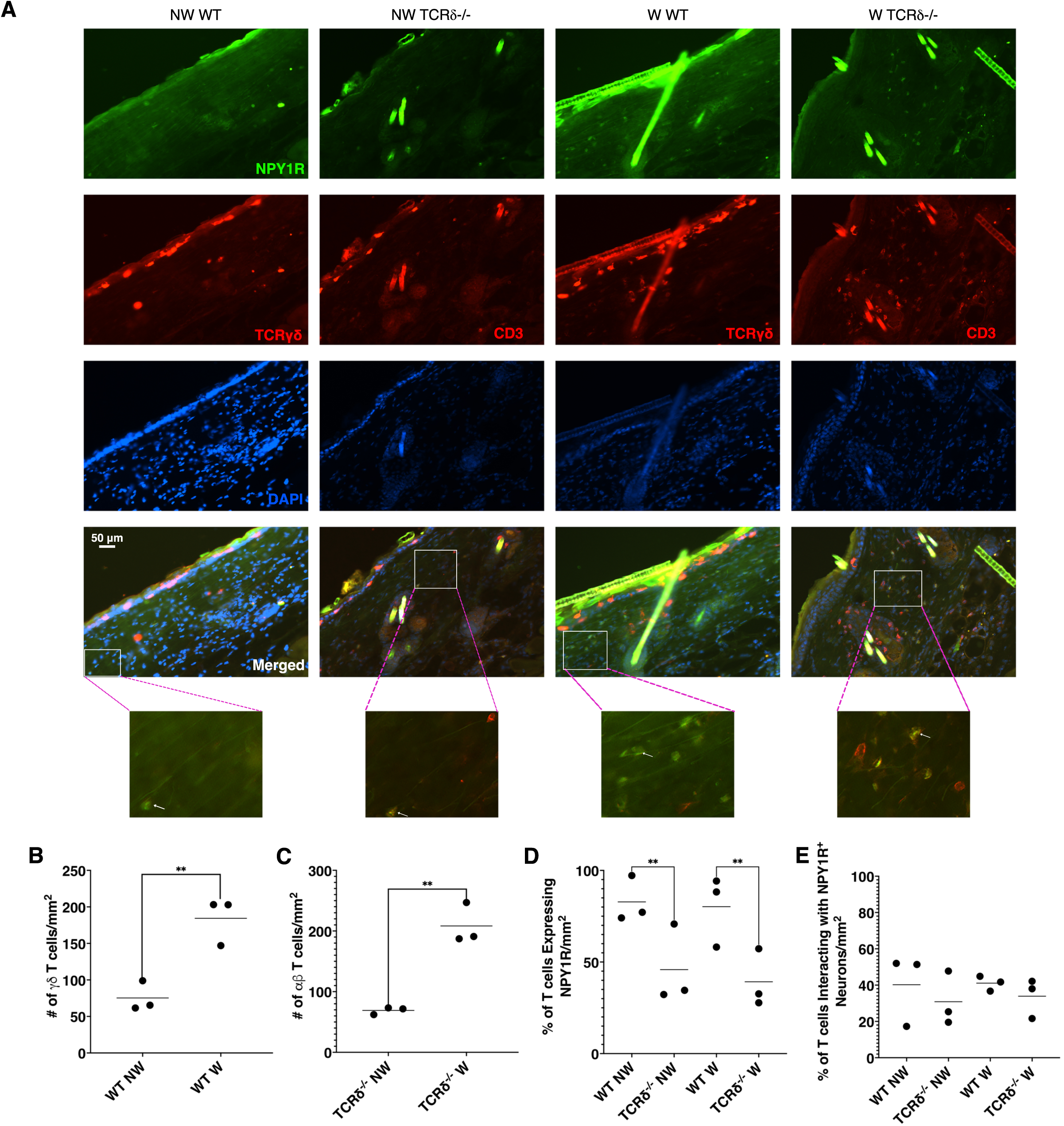
A higher percentage of T cells in the dermis of WT mice express NPY1R compared to T cells in the dermis of TCRδ^−/−^ mice regardless of wound status, while NPY1R^+^ neuron and T cell interactions remain the same. (**A**) Skin sections from non-wounded (NW) or wounded (W) (7 day) WT mice and TCRδ^−/−^ mice were stained with anti-NPY1R FITC (green), anti-TCR γδ PE or anti-CD3 PE (red), and DAPI (blue). (**B**) The number of T cells, (**C**) percentage of T cells expressing NPY1R, and (**D**) percentage of interactions between T cells and NPY1R^+^ neurons were quantified based on fluorescent overlap using immunofluorescence microscopy and Adobe Photoshop (magnification 200X). Two-way ANOVA was used for statistical analysis. \*\**p* < 0.01

### NPY1R is also expressed by γδ T cells in the dermis, but not the epidermis

NPY1R is expressed by immune cells such as splenic lymphocytes, where it inhibits inflammatory cytokine pathways like the TNF signaling pathway (25). However, NPY1R expression by γδ T cells in the skin is not yet studied. We stained epidermal sheets from non-wounded and 10-day wounded WT mice with antibodies specific for NPY1R and Vγ5 to determine if epidermal γδ T cells express NPY1R. We found that epidermal γδ T cells in WT mice do not express NPY1R, regardless of wound status (Fig. 2A). This suggests that epidermal γδ T cells themselves do not require NPY/NPY1R mediated signals at these timepoints.

Next, we examined NPY1R expression by epidermal αβ T cells in TCRδ^−/−^ mice, as certain cytokine receptors essential for the development of epidermal γδ T cells such as CD25, CD69, CD127 and CD215 are reduced on epidermal αβ T cells in TCRδ^−/−^ mice (26). Similar to the WT mice, we found that epidermal αβ T cells do not express NPY1R during nonwounded or wounded conditions (Fig. 2). As dermal and epidermal T cells have distinct roles within the skin, we investigated if dermal T cells express NPY1R. We found that most of the T cells in the dermis express NPY1R both before and after wounding (Fig. 3C). To determine if there is a difference in NPY1R expression by dermal αβ T cells, we stained sections of non-wounded and wounded skin from TCRδ^−/−^ mice with antibodies specific for NPY1R and CD3. We discovered that there is a significantly smaller proportion of T cells expressing NPY1R in TCRδ^−/−^ mice, both before and after wounding (Fig. 3D). This suggests that γδ T cells utilize NPY/NPY1R mediated responses in the dermis. Similar to epidermal γδ T cells in WT mice, we found that epidermal αβ T cells do not express NPY1R during nonwounded or wounded conditions (Fig. 2).

### γδ T cells and NPY1R^+^ neurons interact in the epidermis in a sustained manner during wound repair

Sensory neurons and dermal T cells interact during wound repair, resulting in increased innervation at the wound site (27). Here we investigated epidermal γδ T cell interactions with NPY1R^+^ neurons in WT mice at day 0 and day 10 post-wounding when innervation occurs (Fig. 2A). Approximately half of the epidermal γδ T cells counted interact closely with NPY1R^+^ neurons prior to wounding. Similar numbers of interactions were observed at day 10 post-wounding (Fig. 2B). This data indicates that γδ T cell and NPY1R^+^ neuron interactions are common and sustained.

### Fewer αβ T cell-NPY1R^+^ neuron interactions occur in the epidermis of TCRδ^−/−^ mice but the interactions are sustained during wound repair

We next determined whether epidermal αβ T cells in TCRδ^−/−^ mice also exhibit close interactions with NPY1R^+^ neurons (Fig. 2A). Interactions between αβ T cells and NPY1R^+^ neurons are less common in wounded TCRδ^−/−^ mice than γδ T cells and NPY1R^+^ neurons in wounded WT mice (Fig 2B). There is no significant difference between non-wounded and day 10 wounded T cell-neuron interactions in the TCRδ^−/−^ model (Fig. 2B). Together this indicates that in the epidermis, αβ T cells and NPY1R^+^ neuronal interactions are not impacted by wounding. γδ T cells exhibit a higher tendency to interact with NPY1R^+^ neurons in the epidermis.

### γδ T cells and NPY1R^+^ neurons interact in the dermis in a sustained manner during wound repair

In previous studies examining rosacea-like dermatitis, γδ T cells are in close proximity to calcitonin gene-related peptide (CGRP) expressing neurons and are further recruited to the lesion by CGRP (28). To investigate whether a similar axis exists in the dermis between NPY1R^+^ neurons and γδ T cells, we examined WT mice at day 0 and 7 post-wounding (Fig. 3A). The number of γδ T cells in the dermis is significantly higher on day 7 post-wounding as compared to day 0 (Fig 3B). Approximately 40% of the γδ T cells interact with NPY1R^+^ neurons on both day 0 and 7 post-wounding suggesting that similar to epidermal γδ T cells, dermal γδ T cell-neuron interactions are common and sustained (Fig 3E).

### αβ T cell-NPY1R^+^ neuron interactions occur in the dermis of TCRδ^−/−^ mice and the interactions are sustained during wound repair

We determined whether dermal αβ T cells in TCRδ^−/−^ mice also exhibit close interactions with NPY1R^+^ neurons. The number of αβ T cells increases on day 7 post-wounding as expected due to inflammation (Fig 3C). Approximately 30% of αβ T cells in the dermis interact with NPY1R^+^ neurons on day 0 and 7 post-wounding similar to findings for γδ T cells in WT mice (Fig 3D). This differs with the epidermis, suggesting that γδ T cells exhibit specialized interactions with NPY1R^+^ neurons in the epidermis, while all T cells play roles in communicating with NPY1R^+^ neurons in the dermis.

### Obesity impairs expression of NPY1R in the skin and dermal γδ T cell-NPY1R^+^ neuron interactions

Obesity and type 2 diabetes are associated with elevated levels of TNF-α, which promotes apoptosis of enteric neurons that produce NPY, such as NPY1R^+^ neurons (29, 30). We fed mice either a LFD or HFD for 17 weeks and examined NPY1R expression in the dermis day 0 and day 7 post-wounding. There was a decrease in NPY1R expression in obese mice as compared to lean mice prior to and post-wounding, indicating fewer neurons in the skin and a reduced ability to respond to NPY (Fig 4).

**Figure 4.**
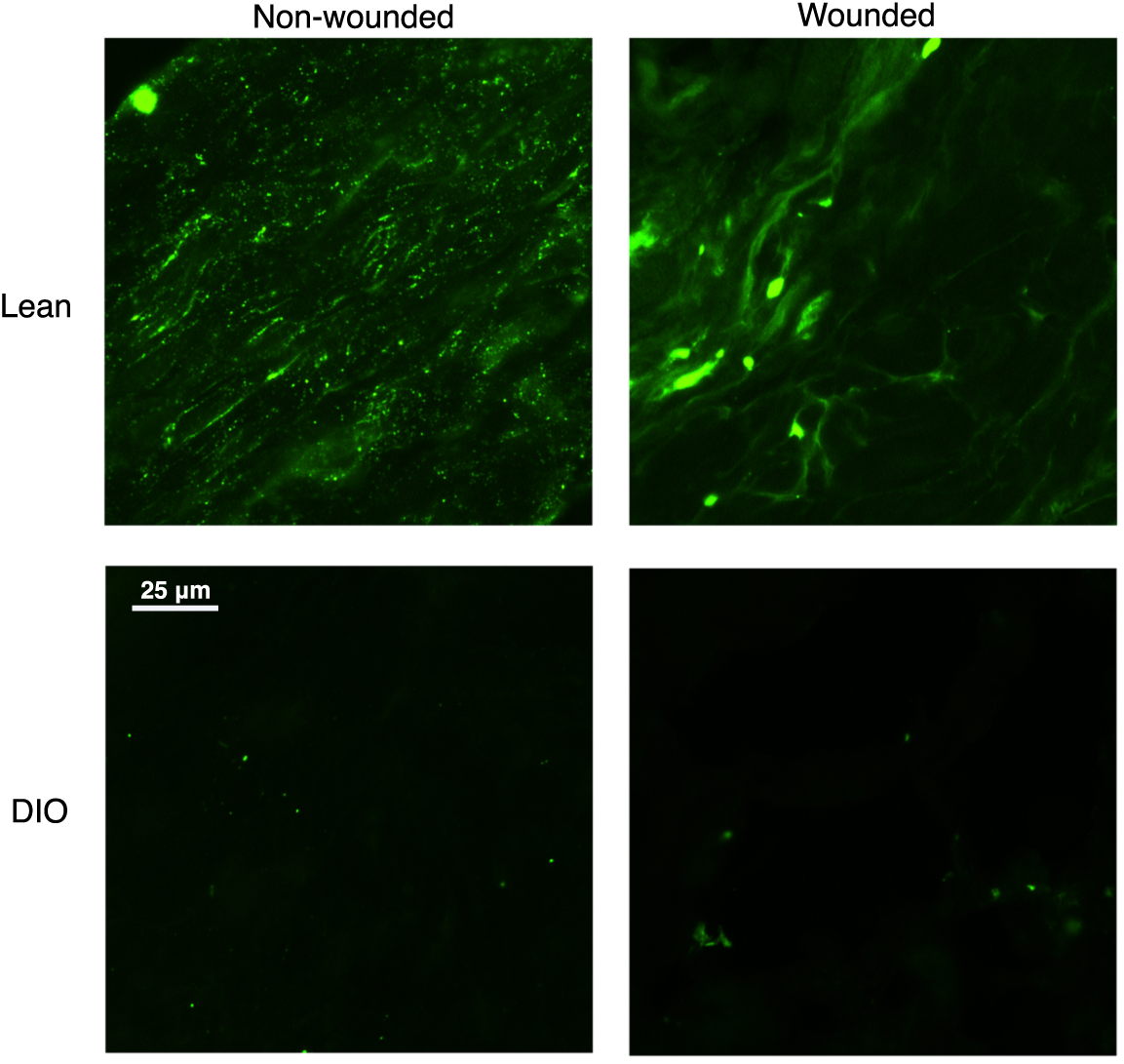
Dermal NPY1R^+^ neurons are reduced in mice fed a HFD as compared to mice fed LFD. Skin sections from lean and obese mice, both non-wounded (NW) and wounded (W), were stained with anti-NPY1R FITC (green) and anti-TCR γδ PE (red). Images were acquired at 200x magnification.

The number of dermal γδ T cells is significantly reduced in the wounded obese mice compared to the wounded lean mice (Fig 5B) similar to previous findings in epidermal γδ T cells (Fig 2B) (21). This suggests that obesity impairs the infiltration of γδ T cells into the wound site. There is a significant decrease in γδ T cell and NPY1R^+^ neurons interactions in obese mice compared to lean mice in both nonwounded and wounded conditions (Fig 5D). However, there is no significant difference in NPY1R-expressing γδ T cells in obese mice as compared to lean mice in non-wounded and wounded conditions (Fig 5C). Overall, the data indicates that diabetes impairs γδ T cell interactions with NPY1R^+^ neurons but not their expression of NPY1R.

**Figure 5.**
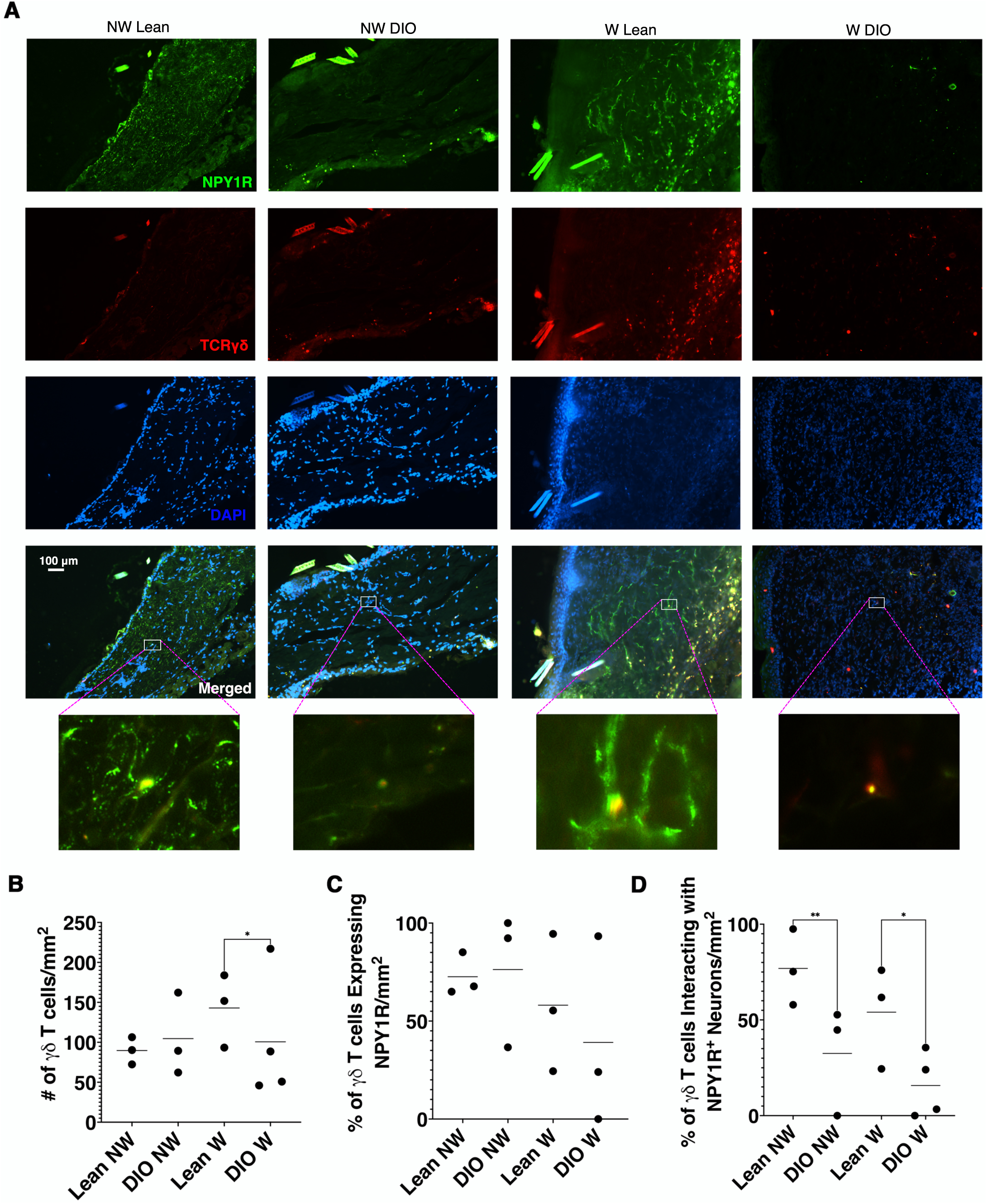
T cells interacting with NPY1R^+^ neurons in the dermis of obese mice are reduced regardless of wound status. (**A**) Skin sections from non-wounded (NW) or wounded (7 day) lean mice and diet-induced obese (DIO) mice were stained with anti-NPY1R FITC (green), anti-TCR γδ PE (red), and DAPI (blue). (**B**) The number of T cells, (**C**) percentage of T cells expressing NPY1R, and (**D**) percentage of interactions between T cells, and NPY1R^+^ neurons were quantified based on fluorescent overlap using immunofluorescence microscopy and Adobe Photoshop (magnification 200X). Two-way ANOVA was used for statistical analysis. \**p* <0.05

## Discussion

γδ T cells participate in neuronal crosstalk in the central nervous system to regulate anxiety-like behavior and short-term memory (31, 32). These γδ T cell-neuronal interactions are regulated by cytokines such as IL-17 and can lead to neuroinflammation (31, 32). Here we analyze interactions between epidermal and dermal γδ T cells and NPY1R-expressing peripheral neurons, which have implications for wound repair and skin inflammation. Following injury, mice lacking γδ T cells do not have a change in baseline sensitivity and pain sensing (33). This suggests that although γδ T cells engage in neuroimmune interactions, their effects may be more regulatory rather than nociceptive. We find that γδ T cells are in close proximity to NPY1R^+^ neurons in nonwounded and wounded skin. Sensory neurons aggravate the pathogenesis of inflammatory skin conditions such as rosacea by the activation of γδ T cell IL-17 production through neuron-produced CGRP (28). Additionally, IL-3-producing epidermal γδ T cells induce higher allergen responses by sensory neurons (24). The ability of neurons and γδ T cell crosstalk to drive neuroinflammation via cytokines suggests close neuroimmune communication that may be important during wound repair.

Epidermal γδ T cells rapidly respond to damaged keratinocytes through the canonical Vγ5Vδ1 TCR with the production of growth factors and cytokines (3, 4, 7, 34, 35). In mice lacking γδ T cells, epidermal γδ T cells are replaced by αβ T cells that lack wound repair responses and express an MHC independent TCR repertoire (26, 36). Our findings that αβ epidermal T cells in TCRδ^−/−^ mice exhibit fewer interactions with neurons in the wound site as compared to epidermal γδ T cells, further substantiates a lack of functional responsiveness by epidermal αβ T cells in wound repair. Additionally, in a murine model of emphysema, NPY knockout mice exhibit elevated IL-17A expression in lung tissue compared to wild type mice suggesting NPY could play an inhibitory role during IL-17A signaling (41). While this has not been directly studied in the skin, a similar mechanism could be at play during cutaneous wound repair. During skin inflammation such as psoriasis or wound healing, IL-17RA is upregulated by neurons (27, 37), allowing them to respond to IL-17A released by IL-17 producers such as epidermal and dermal γδ T cells (9, 38–40). This IL-17A and IL-17RA signaling induces neuronal regeneration (27), but this process likely requires tight regulation to prevent excessive signaling. One possible mechanism is the downregulation of IL-17RA expression by neurons coupled with increased NPY expression to suppress further secretion of IL-17A by γδ T cells. Together, this suggests that NPY could serve as a feedback inhibitor of IL-17A during neuroimmune responses, helping to resolve inflammation and restore homeostasis.

In the spleen, the overexpression of NPY results in decreased levels of cytokines and chemokines, while the knockdown of NPY or NPY1R results in elevated levels of TNFα, IL-6, IL-1β, and CCL20 (25). This suggests that NPY and NPY1R mediated interactions have anti-inflammatory effects. Our studies show that a higher proportion of dermal γδ T cells in WT mice express NPY1R compared to dermal αβ T cells in TCRδ^−/−^ mice. Knowing that NPY1R may play an anti-inflammatory effect on immune cells and that dermal γδ T cells are major producers of IL-17, NPY1R expressed on γδ T cells may act as a mechanism of terminating the inflammatory response. Here we have found that NPY1R is expressed on a majority of the neurons in the skin, suggesting that neurons also utilize NPY for homeostasis and wound repair.

Obesity and type 2 diabetes significantly impair skin immune cells, including the function of skin-resident γδ T cells. Elevated TNF-α levels reduce their ability to secrete cytokines and respond to keratinocyte stimulation (21). In addition to inflammation, the accumulation of reactive oxygen species leads to oxidative stress that further damages neurons and contributes to neuronal apoptosis (30). The combined effects of TNF-α and oxidative stress promote apoptosis of enteric neurons expressing TNFR1 and TNFR2, resulting in reduced levels of NPY (29, 30). NPY^+^ neuron innervation is reduced in adipose tissue of mice fed a HFD, which results in overall decreased levels of NPY (42). Our study demonstrates that obese conditions reduce NPY1R expression by neurons in both non-wounded and wounded conditions. Together, these findings could suggest that a similar mechanism takes place in the skin during obesity and type 2 diabetes, where chronic levels of TNF-α reduce the NPY1R^+^ and NPY^+^ neuron population. This reduction in NPY would subsequently inhibit the anti-inflammatory interactions between NPY and NPY1R expressed by γδ T cells. Future studies to determine the mechanism of communication utilized by γδ T cells and NPY1R-expressing neurons during wound repair may identify novel targets for therapeutics.

## Abbreviations

NPY: Neuropeptide Y
NPY1R: neuropeptide Y receptor Y1
DIO: diet-induced obesity
NW: nonwounded
W: wounded
LFD: low fat diet
HFD: high fat diet
IL-17: interleukin
TNF-α: tumor necrosis factor α
TCR: T cell receptor

